# Effects of antennal segments defects on blood-sucking behavior in Aedes albopictus

**DOI:** 10.1101/2022.09.29.510077

**Authors:** Yiyuan Zhou, Dongyang Deng, Rong Chen, Qian Chen

## Abstract

After mating, female mosquitoes need a blood meal to promote the reproductive process. The blood-feeding transmits extremely harmful infections including malaria, yellow fever, dengue fever, and other arboviruses, making mosquitoes one of the most harmful creatures to human health. The selection and localization of the host by mosquitoes mainly depends on the trace chemical cues emitted by the host into the environment, and the sense of smell is the main way to perceive these trace chemical cues. However, the knowledge of the olfactory mechanism is not enough to meet the needs of mosquito control. Unlike previous studies that focused on the olfactory receptor recognition spectrum to reveal the olfactory mechanism of mosquito host localization. In this paper, we proposed that artificial defects in mosquito antennal flagellomeres may affect their blood-feeding behavior, and through rationally designed behavioral experiments, we found that the 6th and 7th flagellomeres on *Aedes albopictus* antenna are important in the olfactory detection of host searching. Meanwhile, the morphology and distribution of sensilla on each antenna flagellomere were also analyzed and discussed in this paper.

## Introduction

Mosquitoes are one of the greatest public threats to human beings and are even considered the most dangerous animal on earth[1]. After mating, female mosquitoes require blood to complete their oogenesis, the interaction between mosquitoes and their hosts can not only affects humans and livestock, but also transmit harmful diseases, such as malaria, dengue, West Nile fever, lymphatic filariasis, and Zika, during blood feeding[2-5]. At present, spraying insecticides is effective in preventing the spread of mosquito-borne diseases, but the long-term use of chemical insecticides can lead mosquitoes to develop resistance and cause environmental pollution. Therefore, understanding the biological mechanism of selection and host location of mosquitoes and developing a control method to reduce the contact between mosquitoes and hosts are important for mosquito control in the future.

Olfactory cues are still considered to be the main drivers, although it is possible that other types of cues (e.g. visual and thermal) also play a role in the host detection by mosquitoes[6]. Mosquitoes use their ultrasensitive olfactory system to capture these tiny chemical cues and identify the type of host they are feeding on[7, 8]. Therefore, research on the mosquito olfactory system will help to better understand its mechanism and develop green anti-mosquito products that interfere with its olfactory behavior.

However, how the mosquitoes use their olfactory to locate hosts has been an open question. On the one hand, researchers have been trying to find out which odors influence mosquito behavior by stimulating mosquitoes with different odor molecules and compounds through numerous behavioural studies[9-12]. On the other hand, attempts at the cellular and molecular explanations have yielded a great deal of important information[13-17], but the biological mechanism has not been fully explained. This paper attempts to start from another aspect, combining morphology and behavior to find out the functional regions of olfactory organs, to narrow the range of research targets at the cellular and molecular levels.

Antennae are thought to be the most important olfactory organ in mosquitoes since the number of sensilla distributed on the surface accounts for about 90%[18, 19]. In nature, however, like all other animals, mosquitoes may suffer from disasters that result in damage to their antennae. It was also found in lab-bred mosquitoes that some individual mosquitoes did have incomplete antennae, either due to their developmental defects or obvious acquired damage. So, do these mosquitoes with incomplete antennae still have the ability to pinpoint their hosts? Or, whether there is some olfactory protection or compensation mechanism in the 13 antennal flagellomeres that have evolved over a long period, so that when mosquitoes encounter disasters, the partial loss of flagellomeres does not lead to the reduction of their main olfactory perception abilities (such as olfactory localization needed by feeding and reproduction).

*Aedes albopictus* is an important vector control object in China and is the standard material for the efficacy evaluation of mosquito repellent drugs in China. In this study, the role of different antennal flagellomeres in the bloodsucking process of *Aedes albopictus* was studied through behavioral experiments, and the external morphology and distribution of antennal sensilla were observed by scanning electron microscopy. Discussion of results in an attempt to narrow the functional area of olfactory localization in mosquito blood-sucking.

## Materials & Methods

### Mosquito Rearing for Stock Propagation

The colony of *Ae. Albopictus* used in this study was originated from the Center for Disease Control of Hunan Province (China). Mosquitoes were maintained at 28±1°C and 70-80% relative humidity, with a 14:10 h. light/dark photoperiod according to a described protocol[20]. Larvae were fed on yeast powder and adults were maintained on a 10 % sugar solution. For stock propagation, 4-to 5-d old adult female mosquitoes were allowed to take a blood meal from anesthetized mice to lay eggs. The Institutional Reviews Board of The First Affiliated Hospital of Guizhou University of Traditional Chinese Medicine approved all animal procedures (Ethical Application Ref: 20210054).

### Pre-trial for biting assay

Since it is uncertain whether the mosquito will die due to artificial injury when its antennae are cut off, or whether the wound pain will affect its subsequent behavior, pre-trials are needed to determine the impact of antennae cutting surgery on its life and the healing time of the surgical wound.

4 days after eclosion, 20 female mosquitoes from the same batch were selected randomly, 10 of which were left untreated and fed on 10% glucose water alone as a control group. The other 10 were cold-anesthetized for 1 min and cut off the most terminal section of the mosquito antenna (the 13th flagellomere) with Venus scissors (an ophthalmic surgical instrument) under a stereoscope carefully (Figure 1). Mosquitoes with clipped antennae were put back into independent cage and fed with 10% sugar water as a pre-experiment group.

**Figure 1.**
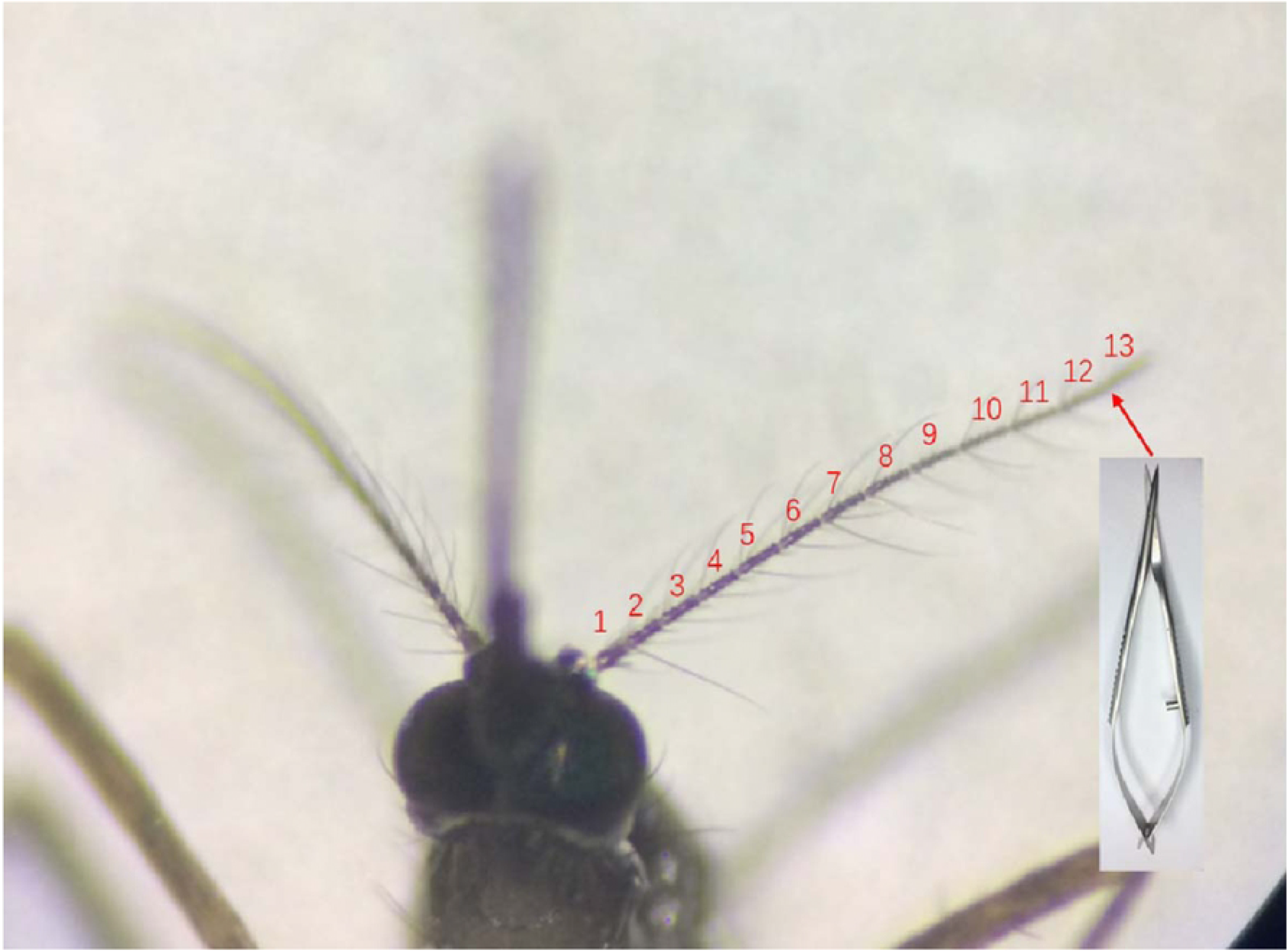
Schematic diagram of pre-experimental mosquitoes. In the pre-experiment, the 13th flagellomere segment at the most distal were cut off with Venus scissors.

Continue to observe and compare the behavior of the mosquitoes in the two cages. 48 hours later, the two groups were deprived of glucose water and fasted for 12 hours. The author put the right hand into the mosquito cage of the control group and the pre-treatment group, respectively, to observe and record the flight and bloodsucking behavior of mosquitoes.

### Biting assay

This assay was based on previously described landing assays[15, 21, 22], and was designed reasonably as follow to reduce errors.

1. The groups. The behavioral experiment in this paper does not cut out each flagellomere of the antennae one by one, but uses a two-step method. First, take 3 flagellomere as a unit, and cut them in sequence to initially lock the range of the flagellomere where the blood-sucking behavior of mosquitoes is significantly reduced. Then, within a locked range of three flagellomere segments, the flagellomere segments were cut off one by one to further identify the segment of the flagellum that caused the decline in mosquito blood-sucking behavior. Therefore, behavioral experiments were grouped by the number of missing flagellomere segments, with 10 experimental samples in each group, and 10 biological replicates were arranged to make the total sample size in each group reach 100. Each replicate was derived from the same batch of pupa-emergent mosquito samples, and the samples were age-consistent between biological replicates. To avoid the possibility of accidental death during the mosquito treatment process, resulting in insufficient final sample size, more than the planned number of mosquitoes in the same batch were selected for antennae shearing, and the living and active mosquitoes were selected for the experiment. Since the blood-sucking behavior of mosquitoes will also be affected at different times of the day, the test sequence between parallel experiments is randomly arranged, that is, parallel experiments are not performed in the order of decreasing or increasing flagellomere segments, but randomly shuffled to reduce the influence of test time on the results.
2. The environment. To ensure that the CO2 concentration in the exhaled breath of the volunteers, and the exposed area of the arm are consistent between parallel experiments, each trial was carried out by the same volunteer, wearing the same clothes, making a mark on the right arm 15 cm away from the fist. Whenever the fist was placed in the cage for each assay, the markings on the arms were just at the mouth of the cage to ensure that the arms in the mosquito cages with the same area of the skin surface in each assay. In addition, a platform was fixed 20cm away from the mosquito cage to place the volunteer’s head, so as to keep the distance between the exhalation and the mosquito cage constant for each assay (Figure 2).

**Figure 2.**
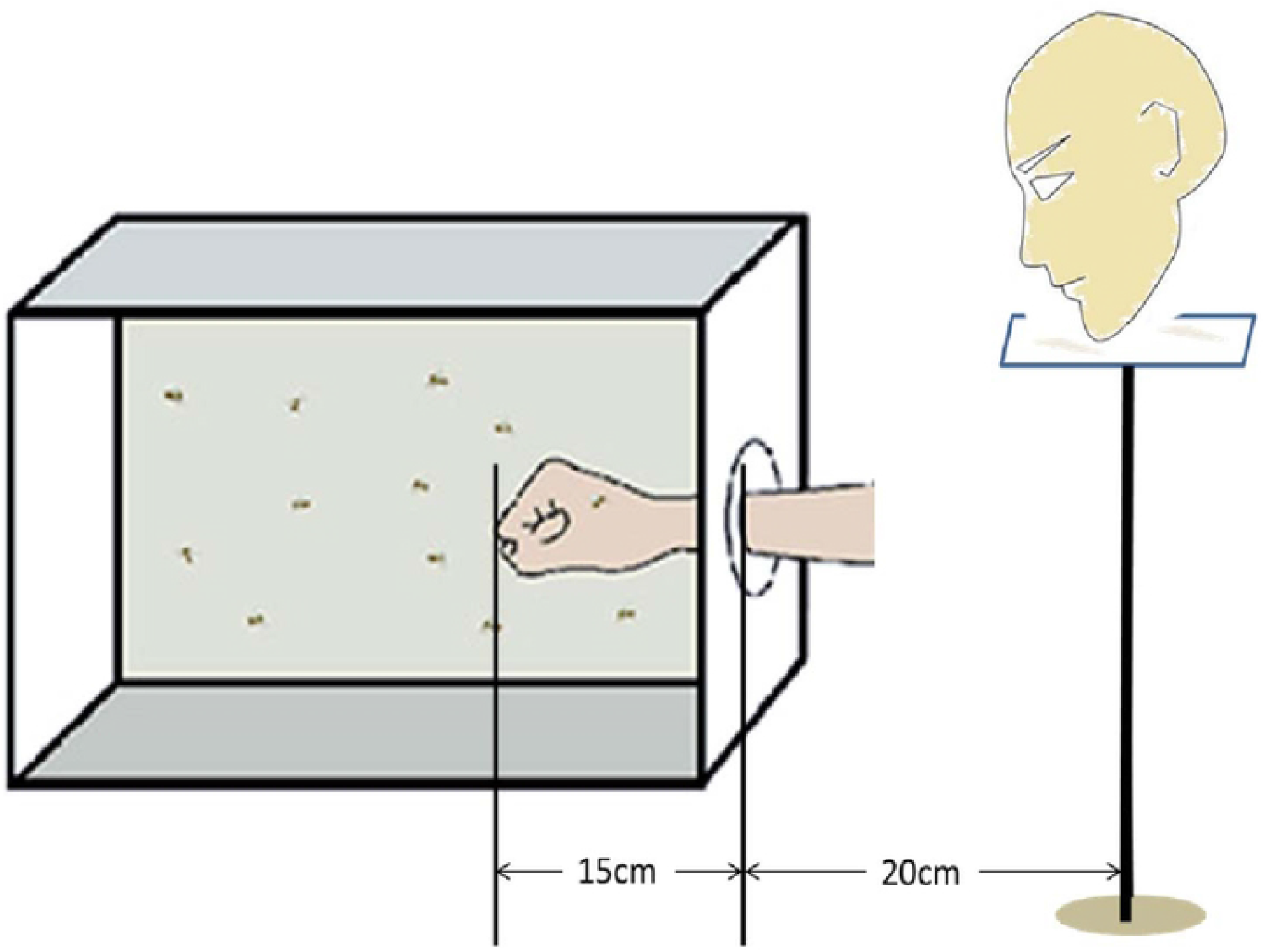
Schematic diagram for behavioral experiment.
3. The volunteer. The volunteer was free from chronic illnesses and did not use any medication regularly, and was requested to avoid garlic, onions, alcohol, or spicy food, take a shower using non-perfumed soap every day, and not participate in strenuous exercise before the test. Half an hour before the test, wash their hands with the same perfume-free soap, and stop the test immediately if feeling unwell.
4. The mosquitoes. The same batch of adult *Ae. albopictus* were reared together and were given free access to 10% sucrose solution. 4 days after eclosion, 12 female mosquitoes were selected randomly for each group from the same batch, and cut the antennal flagellomere segments with Venus scissors under a stereomicroscope carefully after a 1 min cold-anesthesia (Figure 3). The cut female mosquitoes were reared in separate mosquito cages in groups and fed with 10% glucose water for 2 days. Although 12 female mosquitoes treated with antennae were prepared for each group, after fasting for 12 hours, 10 active female mosquitoes were selected for blood-sucking behavior experiment in a random order.

**Figure 3.**
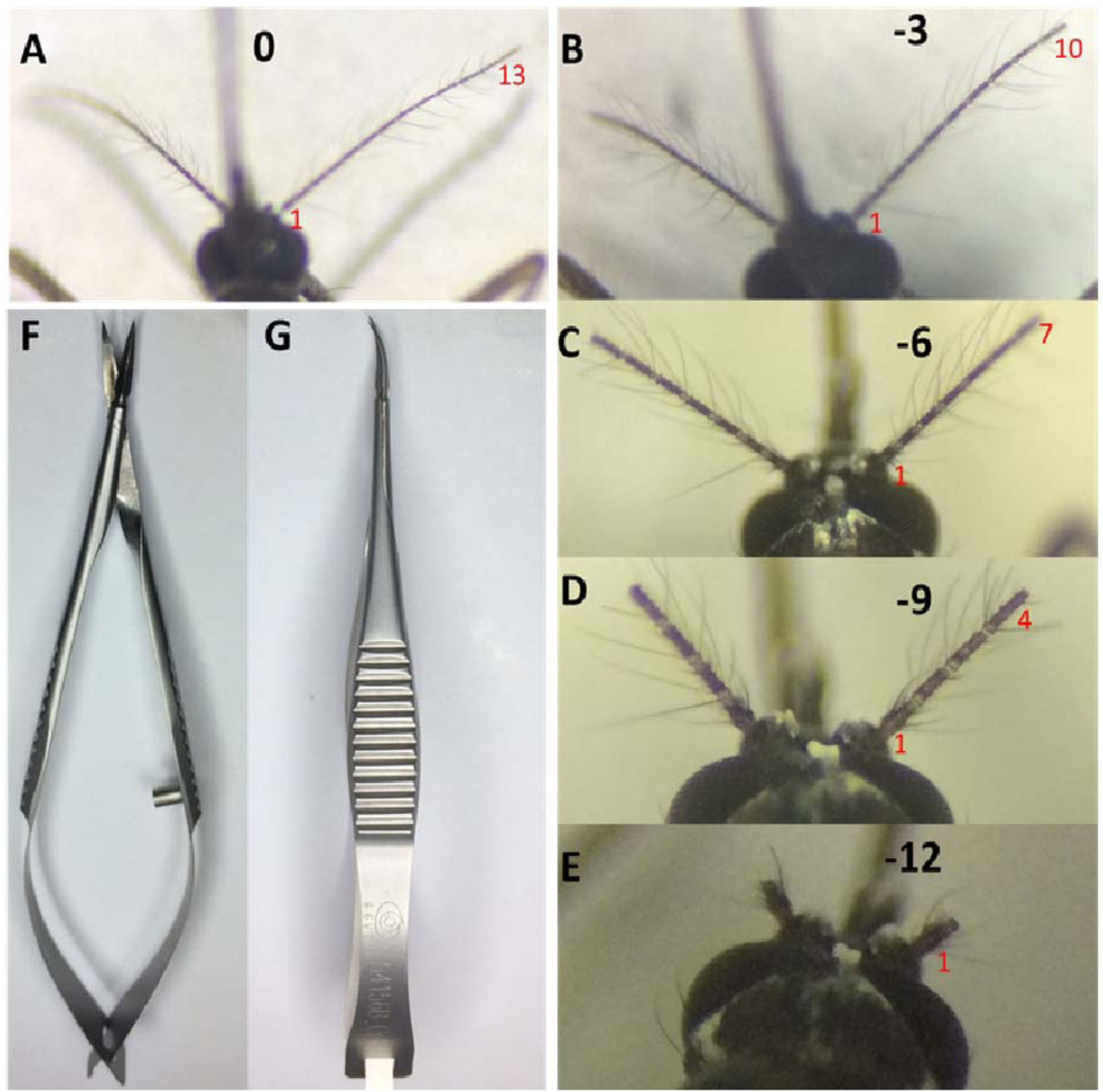
Antennal artificial cut for the first round blood-sucking behavior test. A. Antennal without treatment; B∼E. 3, 6, 9 and 12 segments of antenna were cut off respectively; F∼G. Venus scissors for cutting antennae.
5. The Trial. During the trial, the volunteer (33-year-old female) placed her head on the platform, introduced her arm through cloth sleeves into the cage slowly, and make sure that the mark on her arm was just at the mouth of the cage. Recorded the number of mosquitoes that were blood-fed within 10 min. We defined blood-sucking as landing on the host, inserting the proboscis, and drawing enough blood into the abdomen, or landing on the host, and exploring with the proboscis continuously, which were visible to the naked eye of the observer. After each trial, the volunteers left the chamber for 20 minutes so that the human body odor in the room could be fully dissipated before starting the next trial. Therefore, all trials cannot be completed in one day, but the parallel experiments must be arranged on the same day.

### Scanning electron microscopy (SEM)

Heads with antennae from 4-to 6-day old adult Ae. Albopictus were fixed with 4% paraformaldehyde for 2 h at room temperature, after rinsed with PBS (pH 7.3) containing 0.1% Triton X-100 for 5 times, 50%, 60%, 70%, 80%, 90%, 100% gradient ethanol were used sequential for dehydration. The samples were sequentially rinsed with a mixture of ethanol and hexamethyldisilazane at the ratio of 75:25, 50:50, 25:75, and 0:100, and then air-dried. The dried sample was glued onto aluminum pin mounts with conductive silver, and gold was sprayed on the surface of the sample in a vacuum sprayer. Samples were observed and digital micrographs of each flagellomere were collected using an S4800 scanning electron microscope (Hitachi, Japan).

### Sensilla counts

Sensillae on each flagellomere were classified and counted by morphology. The average for each sensillum type was calculated for 10 individuals and then multiplied by a factor of 2, assuming only half of the sensilla could be seen in each micrograph.

### Statistical analysis

After completion of the biting assays, the percentage of blood-sucking was calculated by dividing the total number of blood-sucking mosquitoes by the total number of mosquitoes used in each bioassay, multiplied by 100. All statistical analyses were performed by GraphPad Prism 8 software (GraphPad Prism), One-way ANOVA and Bonferroni post hoc analysis were used for group comparisons. Data are presented as the mean ± standard error (SEM) (n = 10). Significance was set at p < 0.05.

## Results

### The artificial breakage of the antennae did not cause the death of the mosquito, nor did it affect its activity

The female mosquitoes in the pre-experiment group were all awake after the anesthesia, and there was no death due to antenna breakage. After awakening, mosquitoes will have a short rest (about 20 minutes), and then the flight and feeding behaviors within the next 48 hours were consistent with the control group, showing no listlessness or flight delay. After 12 hours of fasting, mosquitoes in the control group had a strong desire to blood-feed, and the rate of blood-sucking in the control group reached 70% after 10 minutes. While the desire to suck blood was also very strong in the pre-experiment group. Within 2 minutes, 60% of the pre-experiment mosquitoes stayed on the hand and began to suck. After 10 minutes, 50% of the mosquitoes completed blood-feeding, and 30% of the mosquitoes still stayed on the hand to explore and try, the other 20% of the mosquitoes failed to stay on the hand, but they flew for a short time after the hand was introduced into the cage. Although we did not make a statistical analysis, it can be seen that the artificial breakage of the antennae will not cause the death of mosquitoes, nor affect their activities, which was consistent with what we’ve observed in the lab. After 60 hours of operation, the ability of location and sucking were the same as that of untreated mosquitoes. It can be considered that 60 hours after surgery can be used for a follow-up biting assay.

### The 6th and 7th flagellomere of the antennae of Aedes albopictus females may play important roles in their olfactory detection

In the first round with 3 flagellomere segments as the shearing unit, 60 female mosquitoes were divided into 5 groups (Figure 3, A-E), The control group without any treatment (Fig. 3A) and the experimental groups in which 3 (Fig. 3B), 6 (Fig. 3C), 9 (Fig. 3D), and 12 (Fig. 3E) flagellomere segments were cut off from the end of the antennae, respectively. 60 hours after antennae surgery, mosquitoes with 9 and 12 flagellomere segments defection flew briefly in the cage and moved barely since finding a suitable habitat, only a few of them flew back and forth several times before settling on the volunteer’s arm. Through the statistical analysis, the absence of 6 antennae flagellomere segments did not significantly affect the ability of mosquitoes to suck blood, there were still nearly 60% of mosquitoes that can suck blood (Figure 4). However, less than 10% of them suck blood when the antennae were cut by 9 flagellomere segments. That is, the significant difference in the ability of mosquitoes to suck blood occurred between the absence of 6 and 9 antennae flagellomere segments. And it is speculated that the flagellomere segment which affects *Aedes albopictus* to find a host and suck blood may be between or in the 5th and the 7th flagellomere segments.

**Figure 4.**
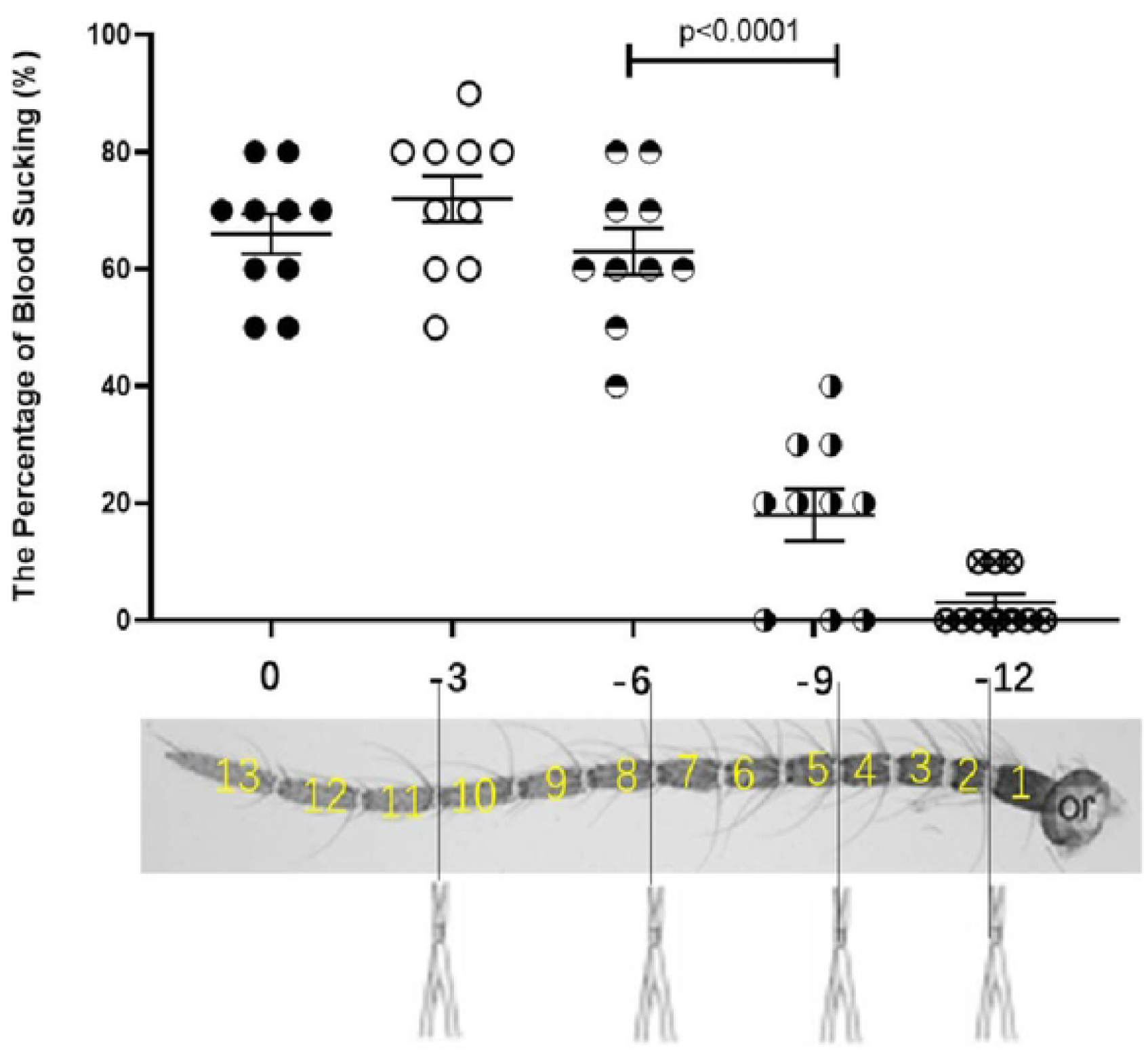
The percentage of blood sucking of Ae. albopictus with antennal segment defection. The horizontal axis represents the number of clipped antennal segments, and the vertical axis represents the average blood sucking percentage. Asterisks indicate significant differences. N=10 groups, 100 mosquitoes in total.

Then, which flagellomere segment plays a key role? In the next round of experiments, the scope will be narrowed, and behavioral testing will be carried out after cutting out flagellomere segments by section.

According to the above method, a total of 84 female mosquitoes were divided into 7 groups, namely: The control group without any treatment and the experimental groups with 3 (Figure 5A), 6 (Figure 5B), 7 (Figure 5C), 8 (Figure 5D), 9 (Figure 5E) and 12 (Figure 5F) flagellomere segments cut off from the end of the antennal, respectively. 60 hours after treatment, behavior difference occurred between the 7 and 8 flagellomere segments defection, nearly half of the mosquitoes had blood-sucking behavior could be shown in 7 flagellomere segments defection, but the flying behavior with 8 flagellomere segments cut off decreased significantly. After a brief period of adaptation, most mosquitoes with 8 flagellomere segments defection did not move until the trial was over, only a small number of them were seen flit around in the cage in each replicate, some of which eventually succeeded in sucking blood (only about 10%), while others stayed on the hand without exploring or feeding (Figure 5G). Therefore, in the segments of 4 to 7 antennae, two segments of antennae flagellomere affect blood-sucking behavior, namely the 6th and the 7th flagellomere. That is, when the mosquito has only 6 flagellate segments left (7 segments defection), the blood-sucking behavior is affected to some extent, but when the mosquito has only 5 flagellate segments left (with 8 segments cut off), the blood-sucking ability was lost almost.

**Figure 5.**
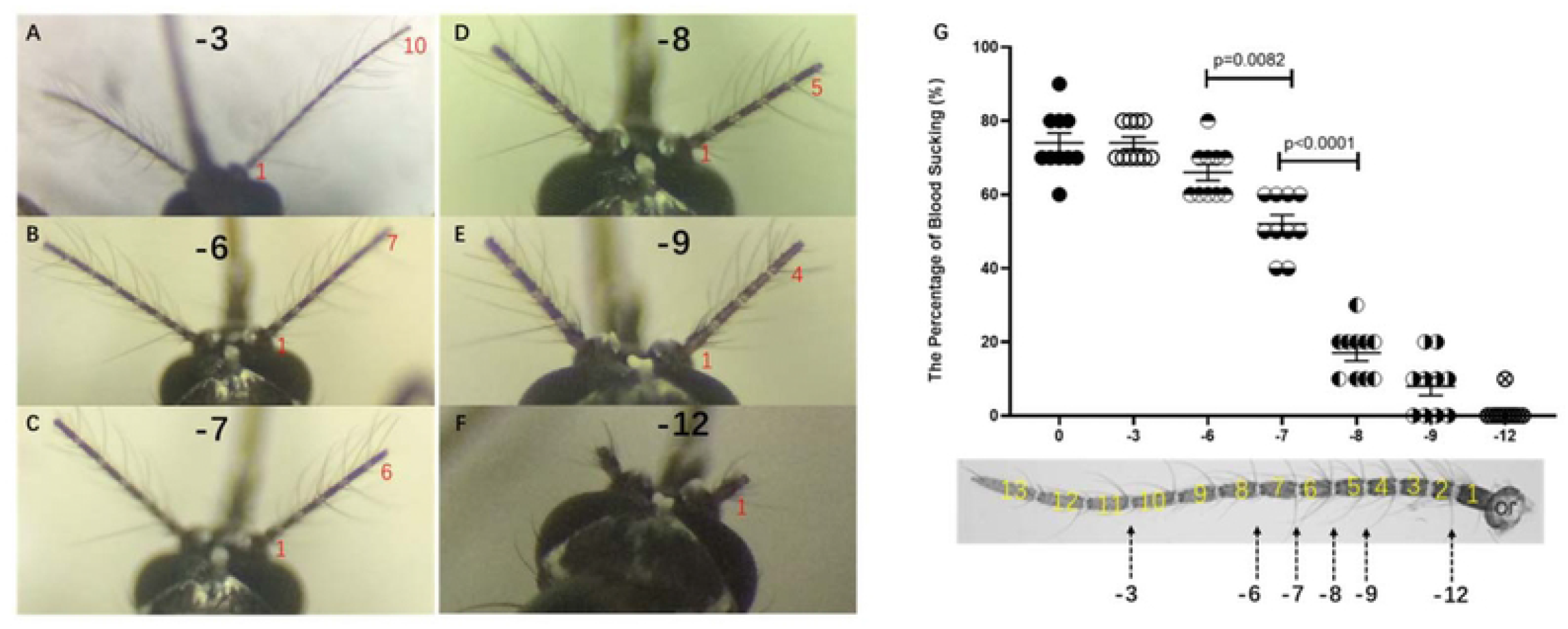
Antennal artificial cut for the second round blood-sucking behavior test. A∼F. 3, 6, 7, 8, 9 and 12 segments of antenna were cut off respectively; G. The percentage of blood sucking of Ae. albopictus with antennal segment defection. N=10 groups, 10 mosquitoes in each group.

### General description of the sensilla of Aedes albopictus

The antennal of *Aedes albopictus* female mosquitoes is divided into 13 distinct flagellomeres with a large number of sensilla. According to the description and nomenclature by Zacharuk[23] and Pitts[24], we classified the sensilla by their morphology (Figure 6). The following are detailed descriptions of the types and distributions of female antennal sensilla.

**Figure 6.**
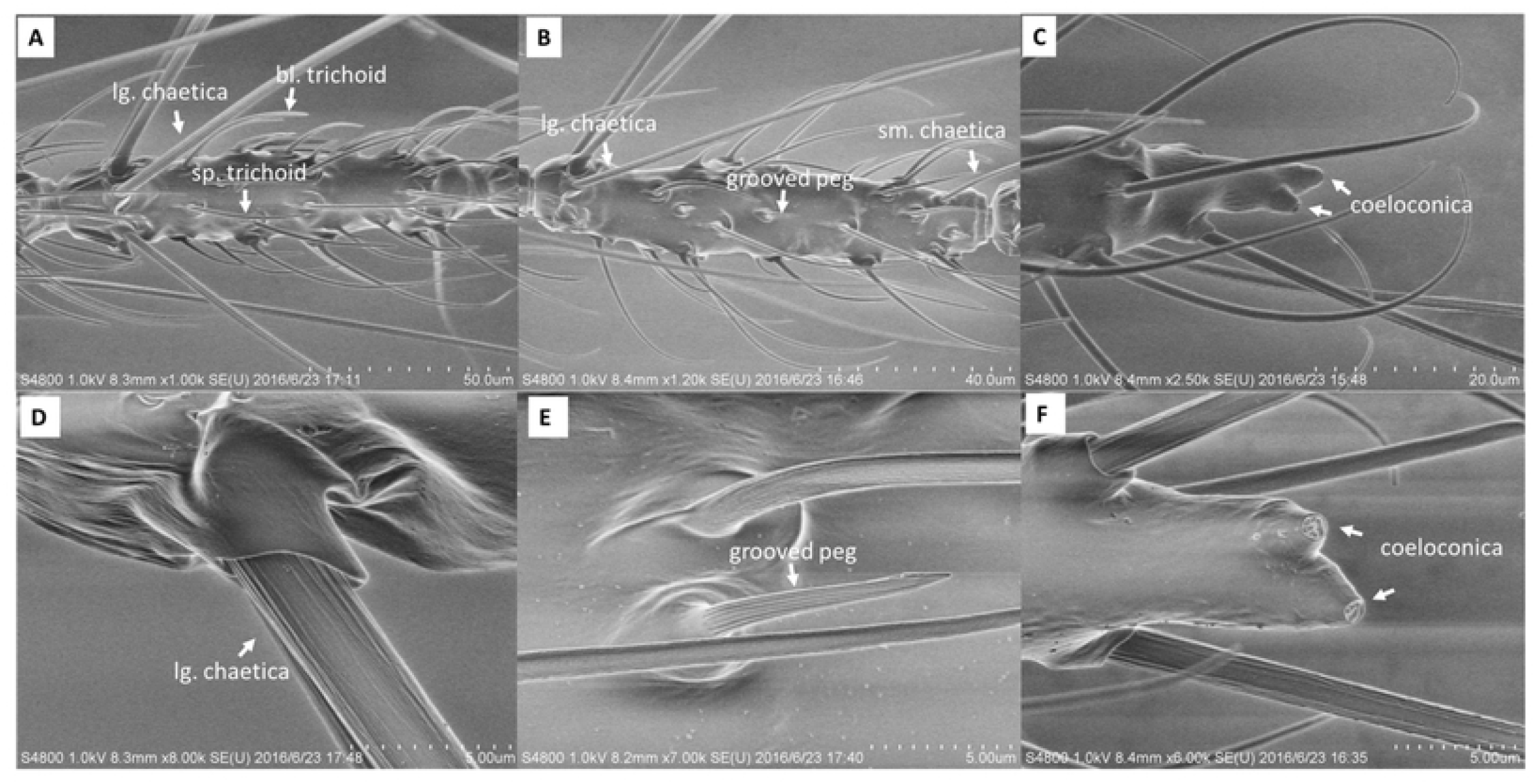
Sensilla types. Representative scanning electron micrographs showing sensilla types found on Ae. albopictus female antennae.

1. Sensilla trichoid: The most widely distributed and most numerous type of sensilla on the antennae with a hair-like structure. According to the shapes, sensilla trichoid were divided into two subtypes: sharp trichodea (sp. trichodea) and blunt trichodea (bl. trichodea) (Figure 6A). sharp trichodea were widely distributed on the antennae, mainly at the 2nd to 13th flagellomeres, and also at the end of the first segment of the flagellum. Blunt trichodea were also mainly distributed at the 2nd to 13th segments of flagellum, but the number is much smaller than that of the sharp ones.
2. Sensilla chaetica: The longest sensilla with grooved and socketed sturdy bristles (Figure 6). Sensilla chaetica were also divided into two subtypes based on length: large chaetica (lg. chaetica) and small chaetica (sm. chaetica) (Figure 6A, B and D). The large chaetica were arranged on the basal end, while the small ones were found nearer the distal edge of flagellomeres 2-13. Their numbers decreased from the proximal to the distal flagellomeres.
3. Sensilla basiconica or grooved peg: The sensilla with grooved but no socketed thorn-shaped hair, which is an important feature that distinguishes them from sensilla chaetica (Fig. 6B and E). The sensilla basiconica were observed on flagellomeres 3–13.
4. Sensilla coeloconica: The smallest sensilla with pitted pegs (Figure 6C and F). The number of such sensilla was also small, they were always found on flagellomeres 1–7, and on the distal tip of the 13th flagellomere with 2.

Finally, considering that there might be a large number of some certain sensilla arranged on flagellomere 6 and 7, which indicated playing important roles in biting assay, the numbers of each sensilla on each flagellomere were used to find some specials (Table 1). However, unfortunately, we saw no evidence of a distinct distribution pattern of different sensilla types on flagellomere 6 and 7.

**Table 1.** types, numbers, and distributions of sensilla (n=10)

| Flagellomere | Sensilla trichoid |  | Sensilla chaetica |  | Sensilla<br>basiconica | Sensilla<br>coeloconica | Total |
| --- | --- | --- | --- | --- | --- | --- | --- |
|  | sp. | bl. | lg. | sm. |  |  |  |
|  | trichodea | trichodea | chaetica | chaetica |  |  |  |
| 1 | 22 | 8 | 10 | 4 | 0 | 3 | 47 |
| 2 | 32.4 | 4.8 | 6 | 6.8 | 1.2 | 0 | 51.2 |
| 3 | 38.4 | 2.4 | 8.8 | 4 | 4.4 | 2 | 60 |
| 4 | 25.2 | 2.4 | 8 | 2.4 | 4 | 0 | 42 |
| 5 | 33.2 | 3.2 | 7.4 | 2 | 3.2 | 3.2 | 52.2 |
| 6 | 27.6 | 3.6 | 6.4 | 2.8 | 4.8 | 1.8 | 47 |
| 7 | 30.2 | 4 | 7.2 | 3.8 | 4.6 | 2 | 51.8 |
| 8 | 31.8 | 4.6 | 6.2 | 3.2 | 4.8 | 0 | 50.6 |
| 9 | 35.2 | 5.8 | 6.2 | 2.2 | 5.2 | 0 | 54.6 |
| 10 | 35.8 | 5.2 | 6.2 | 2.4 | 4.6 | 2.1 | 56.3 |
| 11 | 38.2 | 4.2 | 7 | 3.2 | 5 | 0 | 57.6 |
| 12 | 37.6 | 4.8 | 6.4 | 3.2 | 5.2 | 1.4 | 58.6 |
| 13 | 42.2 | 8.6 | 6.8 | 1 | 8.6 | 4 | 71.2 |
| Total | 429.8 | 61.6 | 92.6 | 41 | 55.6 | 19.4 | 700.1 |

## Discussion and Conclusions

Odor has been demonstrated to be an important cue for host-seeking in mosquitoes, but the underlying biological mechanism remains to be elucidated[15, 25, 26]. The antennae of female mosquitoes are the main peripheral olfactory organs which are composed of 13 flagellomeres, and about 90% of the total number of olfactory sensilla distributed over these antennal subsegments, which made antennae become an important organ in olfactory research. In this study, the hypothesis of the effect of flagellomeres segments on mosquito bloodsucking behavior was proposed, and *Aedes albopictus* was selected as the research object to verify by biting assay. Due to the lack of relevant references, a pre-trial before the biting assay was conducted. The following explanations are required for the pre-trial in this text.

1. About the shearing position of the antennae in the pre-trial. We chose to cut off only the most terminal segment of the mosquito antenna due to the only segment missing has the least impact on its blood-sucking behavior theoretically. Therefore, the blood-sucking behavior can be used to predict the healing time of the antennae. However, if the antennae of mosquitoes were randomly cut off and the pre-trial was conducted, there would be differences in blood-sucking behavior. It is difficult to determine whether these differences in behavior are caused by the absence of the antennal flagellum segment or the unhealed wound.
2. Regarding the determination of time for wound healing. Instead of a series of biting assays at consecutive time points, 60 hours after the operation was selected directly. This is because once the mosquitoes have finished blood-feeding, the next physiological process, digestion and egg laying were started, which led difficult to use the same batch of females for continuous time points biting assays. However, using multiple batches of female mosquitoes to conduct biting assays at different time points requires a large workload. Therefore, we chose to give the mosquitoes a longer healing time, and then conducted the pre-trial directly. The results showed that 60 hours after the operation, there was no behavioral effect due to pain.

In addition, both the mosquito grouping and the test environment were designed in the final biting assays to reduce the experimental error.

In order to understand the attractiveness of humans to the mosquitoes, antennal sensilla have previously been described and compared for several mosquito species. Considering the 6th and 7th flagellomere of the antennae of *Aedes albopictus* females may play a key role in their blood-seeking behavior, it is possible that variations in types or numbers of sensilla exist between them and other flagellomeres, which may suggest areas of future investigation. As such, a comparative examination of the sensilla of each flagellomere on *Aedes albopictus* female antennae was conducted. However, the *Aedes albopictus* females carry the same morphological types of sensilla and the densities of each type are effectively equal between the 6th and 7th flagellomere and other antennal flagellomere. And this result was similar to the result of Qiu’s[27], who identified 6 functional groups of trichoid and 5 functional groups of GP on segments 6–13 of the antenna based on the responses to a panel of 44 compounds in *Anopheles gambiae* by Single-sensillum recordings. The study has shown that the 13th segment distributed the most functional sensilla, which is probably because it is the first part to be exposed to odors, allowing mosquitoes to respond quickly. In the case of the antennae damage, the abundant sensilla on the most distal segment are missing, but the sensilla distributed on other segments of the antennae can be replenished. Qiu’s test indicated that the sensilla on segments 6 and 7 of the antennal flagellum were mainly responsive to some carboxylic acids, alcohols, phenols, ammonia, and amines. Since the authors focussed on segments 6-13 of the antenna, we did not know whether these functional sensilla were also distributed on the segments 1-5.

Based on our findings, we may need to further detect and compare the olfactory-related receptors (including odorant, ionotropic and gustatory receptors) expressed on each segment, which we are trying, albeit with many difficulties.

## References

1. Kamerow D. The world’s deadliest animal. BMJ (Clinical research ed). 2014;348:g3258. Epub 2014/08/19. doi: 10.1136/bmj.g3258. PubMed PMID: 25134129.

2. Guzman MG, Harris E. Dengue. Lancet (London, England). 2015;385(9966):453–65. Epub 2014/09/19. doi: 10.1016/s0140-6736(14)60572-9. PubMed PMID: 25230594.

3. Ashley EA, Pyae Phyo A, Woodrow CJ. Malaria. Lancet (London, England). 2018;391(10130):1608–21. Epub 2018/04/11. doi: 10.1016/s0140-6736(18)30324-6. PubMed PMID: 29631781.

4. Shan C, Xie X, Shi PY. Zika Virus Vaccine: Progress and Challenges. Cell host & microbe. 2018;24(1):12–7. Epub 2018/07/17. doi: 10.1016/j.chom.2018.05.021. PubMed PMID: 30008291; PubMed Central PMCID: PMCPMC6112613.

5. Chandrasegaran K, Lahondère C, Escobar LE, Vinauger C. Linking Mosquito Ecology, Traits, Behavior, and Disease Transmission. Trends in parasitology. 2020;36(4):393–403. Epub 2020/03/20. doi: 10.1016/j.pt.2020.02.001. PubMed PMID: 32191853.

6. Vinauger C, Van Breugel F, Locke LT, Tobin KKS, Dickinson MH, Fairhall AL, et al. Visual-Olfactory Integration in the Human Disease Vector Mosquito Aedes aegypti. Current biology : CB. 2019;29(15):2509-16.e5. Epub 2019/07/23. doi: 10.1016/j.cub.2019.06.043. PubMed PMID: 31327719; PubMed Central PMCID: PMCPMC6771019.

7. Yan J, Gangoso L, Ruiz S, Soriguer R, Figuerola J, Martínez-de la Puente J. Understanding host utilization by mosquitoes: determinants, challenges and future directions. Biological reviews of the Cambridge Philosophical Society. 2021;96(4):1367–85. Epub 2021/03/10. doi: 10.1111/brv.12706. PubMed PMID: 33686781.

8. Takken W, Verhulst NO. Host preferences of blood-feeding mosquitoes. Annual review of entomology. 2013;58:433–53. Epub 2012/10/02. doi: 10.1146/annurev-ento-120811-153618. PubMed PMID: 23020619.

9. Logan JG, Birkett MA, Clark SJ, Powers S, Seal NJ, Wadhams LJ, et al. Identification of human-derived volatile chemicals that interfere with attraction of Aedes aegypti mosquitoes. Journal of chemical ecology. 2008;34(3):308–22. Epub 2008/03/01. doi: 10.1007/s10886-008-9436-0. PubMed PMID: 18306972.

10. Logan JG, Stanczyk NM, Hassanali A, Kemei J, Santana AE, Ribeiro KA, et al. Arm-in-cage testing of natural human-derived mosquito repellents. Malaria journal. 2010;9:239. Epub 2010/08/24. doi: 10.1186/1475-2875-9-239. PubMed PMID: 20727149; PubMed Central PMCID: PMCPMC2931528.

11. Bernier UR, Kline DL, Barnard DR, Schreck CE, Yost RA. Analysis of human skin emanations by gas chromatography/mass spectrometry. 2. Identification of volatile compounds that are candidate attractants for the yellow fever mosquito (Aedes aegypti). Analytical chemistry. 2000;72(4):747–56. Epub 2000/03/04. doi: 10.1021/ac990963k. PubMed PMID: 10701259.

12. Bernier UR, Kline DL, Schreck CE, Yost RA, Barnard DR. Chemical analysis of human skin emanations: comparison of volatiles from humans that differ in attraction of Aedes aegypti (Diptera: Culicidae). J Am Mosq Control Assoc. 2002;18(3):186–95. Epub 2002/09/27. PubMed PMID: 12322940.

13. Kwon HW, Lu T, Rützler M, Zwiebel LJ. Olfactory responses in a gustatory organ of the malaria vector mosquito Anopheles gambiae. Proceedings of the National Academy of Sciences of the United States of America. 2006;103(36):13526–31. Epub 2006/08/30. doi: 10.1073/pnas.0601107103. PubMed PMID: 16938890; PubMed Central PMCID: PMCPMC1569196.

14. Leal WS. Odorant reception in insects: roles of receptors, binding proteins, and degrading enzymes. Annual review of entomology. 2013;58:373–91. Epub 2012/10/02. doi: 10.1146/annurev-ento-120811-153635. PubMed PMID: 23020622.

15. McBride CS, Baier F, Omondi AB, Spitzer SA, Lutomiah J, Sang R, et al. Evolution of mosquito preference for humans linked to an odorant receptor. Nature. 2014;515(7526):222–7. Epub 2014/11/14. doi: 10.1038/nature13964. PubMed PMID: 25391959; PubMed Central PMCID: PMCPMC4286346.

16. Chen Q, Man Y, Li J, Pei D, Wu W. Olfactory Ionotropic Receptors in Mosquito Aedes albopictus (Diptera: Culicidae). Journal of medical entomology. 2017;54(5):1229–35. Epub 2017/04/12. doi: 10.1093/jme/tjx063. PubMed PMID: 28399284.

17. Pitts RJ, Derryberry SL, Zhang Z, Zwiebel LJ. Variant Ionotropic Receptors in the Malaria Vector Mosquito Anopheles gambiae Tuned to Amines and Carboxylic Acids. Scientific reports. 2017;7:40297. Epub 2017/01/10. doi: 10.1038/srep40297. PubMed PMID: 28067294; PubMed Central PMCID: PMCPMC5220300.

18. McIver SB. Sensilla mosquitoes (Diptera: Culicidae). Journal of medical entomology. 1982;19(5):489–535. Epub 1982/10/14. doi: 10.1093/jmedent/19.5.489. PubMed PMID: 6128422.

19. Isberg E, Hillbur Y, Ignell R. Comparative study of antennal and maxillary palp olfactory sensilla of female biting midges (Diptera: Ceratopogonidae: Culicoides) in the context of host preference and phylogeny. Journal of medical entomology. 2013;50(3):485–92. Epub 2013/06/28. doi: 10.1603/me12235. PubMed PMID: 23802442.

20. Chen XG, Jiang X, Gu J, Xu M, Wu Y, Deng Y, et al. Genome sequence of the Asian Tiger mosquito, Aedes albopictus, reveals insights into its biology, genetics, and evolution. Proceedings of the National Academy of Sciences of the United States of America. 2015;112(44):E5907–15. Epub 2015/10/21. doi: 10.1073/pnas.1516410112. PubMed PMID: 26483478; PubMed Central PMCID: PMCPMC4640774.

21. Smallegange RC, van Gemert GJ, van de Vegte-Bolmer M, Gezan S, Takken W, Sauerwein RW, et al. Malaria infected mosquitoes express enhanced attraction to human odor. PloS one. 2013;8(5):e63602. Epub 2013/05/22. doi: 10.1371/journal.pone.0063602. PubMed PMID: 23691073; PubMed Central PMCID: PMCPMC3655188.

22. Verhulst NO, Qiu YT, Beijleveld H, Maliepaard C, Knights D, Schulz S, et al. Composition of human skin microbiota affects attractiveness to malaria mosquitoes. PloS one. 2011;6(12):e28991. Epub 2012/01/05. doi: 10.1371/journal.pone.0028991. PubMed PMID: 22216154; PubMed Central PMCID: PMCPMC3247224.

23. Zacharuk RY, Shields VD. Sensilla of Immature Insects. Annual review of entomology. 1991;36(1):331–54.

24. Pitts RJ, Zwiebel LJ. Antennal sensilla of two female anopheline sibling species with differing host ranges. Malaria journal. 2006;5:26. Epub 2006/04/01. doi: 10.1186/1475-2875-5-26. PubMed PMID: 16573828; PubMed Central PMCID: PMCPMC1532926.

25. Wang G, Carey AF, Carlson JR, Zwiebel LJ. Molecular basis of odor coding in the malaria vector mosquito Anopheles gambiae. Proceedings of the National Academy of Sciences of the United States of America. 2010;107(9):4418–23. Epub 2010/02/18. doi: 10.1073/pnas.0913392107. PubMed PMID: 20160092; PubMed Central PMCID: PMCPMC2840125.

26. Carey AF, Wang G, Su CY, Zwiebel LJ, Carlson JR. Odorant reception in the malaria mosquito Anopheles gambiae. Nature. 2010;464(7285):66–71. Epub 2010/02/05. doi: 10.1038/nature08834. PubMed PMID: 20130575; PubMed Central PMCID: PMCPMC2833235.

27. Qiu YT, van Loon JJ, Takken W, Meijerink J, Smid HM. Olfactory Coding in Antennal Neurons of the Malaria Mosquito, Anopheles gambiae. Chemical senses. 2006;31(9):845–63. Epub 2006/09/12. doi: 10.1093/chemse/bjl027. PubMed PMID: 16963500.

